# Developmental Trajectories of Addition and Subtraction in Early School-aged Children

**DOI:** 10.1101/2025.11.27.691071

**Authors:** Manqi Zhou, Shiyan Ji, Yalun Zhang, Huijuan Shen, Jipeng Huang, Qinfen Zhang, Yiwen Li, Haitian Mei, Rong Kuang, Yuanxin Lin, Yan Song, Jing Huang, Xuan Dong

## Abstract

Mastering basic arithmetic lays the foundation for lifelong mathematical achievement, yet how the neural mechanisms underlying addition and subtraction calculations develop during childhood remain unclear. This study investigated the developmental trajectories of addition and subtraction calculations in school-aged children, and explored how arithmetic-specific neural responses in the right parieto-occipital region are related to age-related improvements in behavioral performance. Here, we recorded electroencephalography (EEG) signals from 115 typically developing children divided into three age groups (7–8-, 9–10-, and 11–12-year-olds) during an arithmetic task involving addition and subtraction calculations. We analyzed the P300 component from the original waveforms and the ΔP300 component from the difference waveforms, which were generated by subtracting the event-related potentials (ERPs) elicited by the first operand from those elicited by the second operand to isolate arithmetic-specific neural processes. The results revealed that, in both calculations, older children showed better behavioral performance (faster reaction time and larger PI) and greater right parieto-occipital ΔP300 amplitudes, with a key developmental stage at 11–12 years. Furthermore, all groups of children exhibited greater right parieto-occipital P300 and ΔP300 amplitudes in subtraction, whereas the smaller right parieto-occipital ΔP300 amplitudes in addition significantly mediated the relationship between age and PI. These findings reveal distinct patterns for addition and subtraction calculations over development, highlighting the right parieto-occipital ΔP300 amplitude as a neural marker of arithmetic performance, particularly for addition calculation.

## 1. Introduction

Mathematical abilities develop primarily during childhood and are applied throughout daily life. Mastering basic arithmetic forms the foundation for more advanced mathematical learning and relies on some general cognitive processes, such as working memory, attention, and verbal skills (Anderson, 2000; Butterworth, 2005). Given the central role of arithmetic in both education and daily functioning, understanding how the brain supports these skills has become a major focus in developmental cognitive neuroscience. In particular, identifying the neural mechanisms involved in arithmetic processing helps inform educational practices and detect early developmental disorders (Butterworth et al., 2011; Menon, 2010).

Building on this perspective, previous research has used a delayed answer verification task to investigate the neural processing related to basic arithmetic (De Smedt et al., 2009; Jasinski and Coch, 2012; Zhou et al., 2006). The positive component peaking at approximately 300 ms (P300), associated with arithmetic fact retrieval (Muluh et al., 2011; Zhou et al., 2006) and complexity of cognitive processing (Gao et al., 2022), has been served as a marker of pre-solution cognitive processing differences between arithmetic operations (Jost et al., 2004). Further studies reported that the P300 amplitude at central and parietal electrodes was significantly larger for subtraction compared with addition (Muluh et al., 2011; Núñez-Peña et al., 2004). Moreover, neuroimaging studies revealed greater activation for subtraction than for addition in the posterior parietal cortex, including the bilateral intraparietal sulcus (IPS) (Dehaene et al., 2003; Ischebeck et al., 2009; Menon, 2010; Prado et al., 2011). These findings suggest distinct cognitive mechanisms between addition and subtraction, underscoring the importance of understanding how arithmetical skills mature in the brain and how such insights can inform strategies to support mathematical learning in educational settings.

Further evidence also suggests that addition and subtraction activate distinct patterns in the left and right hemispheres. Based on adult data, the triple-code model posits that addition relies primarily on a left-lateralized network that supports direct access to verbal memory for fact retrieval, whereas subtraction depends on a bilateral system that supports quantity-based manipulations for magnitude representation (Dehaene and Cohen, 1997). Specifically, the right parieto-occipital cortex, such as the IPS, is a critical region of this bilateral network in children, supporting fundamental processes like magnitude representation and visuospatial working memory (Price et al., 2007). During subtraction tasks, children showed pronounced activation in the right IPS, reflecting nonverbal quantity processing (Chochon et al., 1999; Kawashima et al., 2004). During addition tasks, the right IPS is also crucial for processing spatial-numerical associations, such as the sequential representation of numbers along the mental number line, thereby influencing arithmetic performance (Montefinese et al., 2017; Salillas et al., 2023). These findings highlight distinct functional roles of the right parieto-occipital region in addition and subtraction calculations. Although previous evidence has observed that activation in the right parietal region increases with age during arithmetic tasks (Evans et al., 2016), it remains unclear whether the involvement of the right parieto-occipital region differs between addition and subtraction, and how it relates to developmental improvements in arithmetic performance.

Arithmetic processing during childhood is closely linked to brain maturation. As children develop, more efficient allocation of cognitive resource is associated with better arithmetic performance (Anderson, 2000). Research has further revealed that age influences mathematical cognition indirectly through verbal working memory and processing speed (Formoso et al., 2018). Rosenberg-Lee et al. (2011) reported that children from second to third grade showed increased activation in right superior parietal and lateral occipital cortex, reflecting greater integration among attention, working memory, and numerical processing systems. Given the importance of attention and working memory in arithmetic calculation, developmental changes of arithmetic processing may be reflected in the P300 component, which is sensitive to these cognitive functions (Polich, 2007). Dong et al. (2007) revealed that P300 latencies decreased with age in 8-, 9-, and 11-year-old children when they performed arithmetic tasks, suggesting increased neural efficiency in arithmetic processing with age. However, it remains unclear whether, and to what extent, the development of arithmetic performance is directly driven by arithmetic-specific neural mechanisms. Therefore, further investigations of the neural and behavioral correlates of arithmetic development are needed (Kaufmann et al., 2011). To address this, we examined whether developmental improvements in behavioral performance were associated with arithmetic-specific neural responses in the right parieto-occipital region, based on prior evidence highlighting this region in magnitude manipulations during subtraction and in spatial-numerical associations during addition.

Given the paucity of the extant research, we aim to investigate the development trajectories of addition and subtraction calculations in school-aged children. Most previous studies have simultaneously presented an arithmetic problem or solution (e.g., a + b = or a + b = c), focusing on the ERPs time-locked to the presentation of the problem or answer. However, presenting a whole equation involves multiple general cognitive processes, such as the retention of two operands, processing of the operator sign, and maintenance of intermediate results. Here, we adopted a new paradigm by presenting each item sequentially to minimize these general cognitive processes, and derived a ΔP300 component from the difference waves of the two operands to isolate arithmetic-specific neural processes. We hypothesize that both the right parieto-occipital P300 and ΔP300 amplitudes will be greater for subtraction than for addition, because subtraction requires greater cognitive resources for working memory demands and numerical magnitude manipulations (Chassy and Grodd, 2012; Holloway and Ansari, 2010). Moreover, older children may exhibit greater right parieto-occipital ΔP300 amplitudes and perform both calculations better, indicating the maturation of neural mechanisms that support greater arithmetic performance. By analyzing P300 and ΔP300 amplitudes across three groups of children (7–8-, 9–10-, and 11–12-year-olds), we shed novel light on a more comprehensive description of the neural maturation of arithmetic processing during childhood.

## 2. Methods

### 2.1. Participants

The sample size was estimated by conducting a prior power analysis with G*power (Faul et al., 2007) within a repeated-measures analysis of variance (ANOVA) with two repeated measurements and three groups. The results indicated the required total sample size to be 105 participants for within-between interactions, with a medium effect size of *f* = .25, a power of 90%, and an alpha level of .05. On the basis of this power analysis, and to account for a possible data dropout, the data collection was set out to obtain data from 115 Chinese children, in which 10 children were excluded due to either too few trials (3 children with low behavioral performance and 5 children with poor electroencephalogram [EEG] quality) or technical issues (2 children). Thus, a total of 105 children were included for the final analyses in this study and were categorized into three age groups: 7–8-year-olds (*n* = 29, 5 girls and 24 boys, mean age = 8.19 ± 0.54), 9–10-year-olds (*n* = 36, 12 girls and 23 boys, mean age = 9.99 ± 0.57), and 11–12-year-olds (*n* = 40, 12 girls and 28 boys, mean age = 11.92 ± 0.45).

All participants were right-handed, and had normal or corrected-to-normal vision. None had a history of neurological or psychiatric disorders, severe developmental diseases, or current use of psychoactive medications. Each participant had a full-scale intelligence quotient (IQ) score ≥ 80, as assessed by the validated version of the Wechsler Intelligence Scale for Children-Revised (WISC-R; Wechsler, 1974). The participants’ demographic characteristics for the three age groups are presented in Table 1. Age differed significantly across the three age groups (*F*_(2,102)_ = 441.91, *p* < 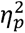 = 0.897). No significant group differences were found in sex distribution (χ^2^ = 2.26, *p* = .322, φ = 0.147) or full-scale IQ (*F*_(2,102)_ = 0.12, *p* = .884, rJ^2^ = 0.002). Both verbal and written informed consent was obtained from all of the children and their parents prior to the experiment, in accordance with the protocols approved by the local Ethics Committee.

**Table 1.** Demographic information including sex, age and full-scale IQ score per age group.

| | 7–8-year-olds<br>($n = 29$) | 9–10-year-olds<br>($n = 36$) | 11–12-year-olds<br>($n = 40$) | Group differences | | |
| --- | --- | --- | --- | --- | --- | --- |
| | | | | $F / \chi^2$ | $p$-value | $\eta_p^2 / \phi$ |
| Sex (girls: boys) | 5: 24 | 12: 24 | 12: 28 | 2.26 | .322 | 0.147 |
| Age in years | $8.19 \pm 0.54$ | $9.99 \pm 0.57$ | $11.92 \pm 0.45$ | 441.91 | <.001*** | 0.897 |
| Full-scale IQ | $102.21 \pm 14.67$ | $102.81 \pm 12.28$ | $102.37 \pm 13.47$ | 0.12 | .884 | 0.002 |

*Note.* Data are presented as the mean ± standard deviation (SD), unless otherwise indicated. $\chi^2$ = chi-square statistic, $n$ = number of children included in each age group, IQ = intelligence quotient. \*\*\* $p < .001$ .

### 2.2. Stimuli and procedure

Participants were seated comfortably in a chair 70 cm away from a 19-inch monitor in a dimly lit, sound-attenuating and electrically-shielded chamber. They first completed 8 practice trials and received feedback after each response. Then, during the recording period, they did not receive any feedback regarding accuracy for each trial to avoid demotivation. They performed an arithmetic task including addition and subtraction calculations, with all numbers ranging from 2 to 20. Problems with 0 or 1 as an operand (e.g., 6 + 0 or 6 – 1) and those with same operands (e.g., 5 + 5 or 5 – 5) were excluded from both calculations, because they are rule-based arithmetic facts (LeFevre and Liu, 1997) or tend to elicit atypical behavioral responses (Blankenberger, 2001; Campbell and Graham, 1985). For all the calculations, half of the proposed solutions were congruous with the correct answer, whereas half were incongruous. To account for the distance effect, incongruous solutions were generated by adding or subtracting 1 or 2 from the correct answer (e.g., congruous: 3 + 4 = 7, incongruous: 3 + 4 = 9). Addition calculations without carrying or subtraction calculations without borrowing were defined as simple problems, whereas addition calculations with carry or subtraction calculations with borrow were defined as complex problems (Kong et al., 2005). There was no significant difference in the numbers of simple and complex problems between addition and subtraction (*t*_(100)_ = 0.41, *p* = .686, Cohen’s *d* = 0.082), suggesting that two calculations were comparable in the level of difficulty. To mitigate artifacts from blinking and eye movements, the participants were encouraged to keep their eyes constantly on the screen center and remain still to limit muscle and ocular artifacts.

The experiment was programmed using E-prime software (Psychology Software Tools Inc., Pittsburgh, PA, USA). All stimuli were presented in white text against a black background (Fig. 1A). Each trial started with the presentation of a fixation point for 600 ms, followed by a delay of 500 ms. The operands appeared sequentially in the screen center as Arabic numerals, with an operator symbol (“+” or “–”) displayed between them. Each of them lasted for 300 ms and the intervals were 500 ms. A blank screen was presented for 2000 ms after the second operand vanished. The solution remained visible on the screen until the participant responded or for a maximum period of 2000 ms. Participants were instructed to justify whether the solution was congruous or incongruous with the correct answer and to respond as quickly and accurately as possible. The next trial began after a response or a maximum period of 1500 ms. The order of trials was counterbalanced, with the constraint that no more than two congruous or incongruous solutions were presented in a row, and that consecutive problems did not have the same operand or solution. The task lasted ∼30 minutes and consisted of 102 trials with eight practice trials to ensure that the participants understand the task instructions.

**Fig. 1.**
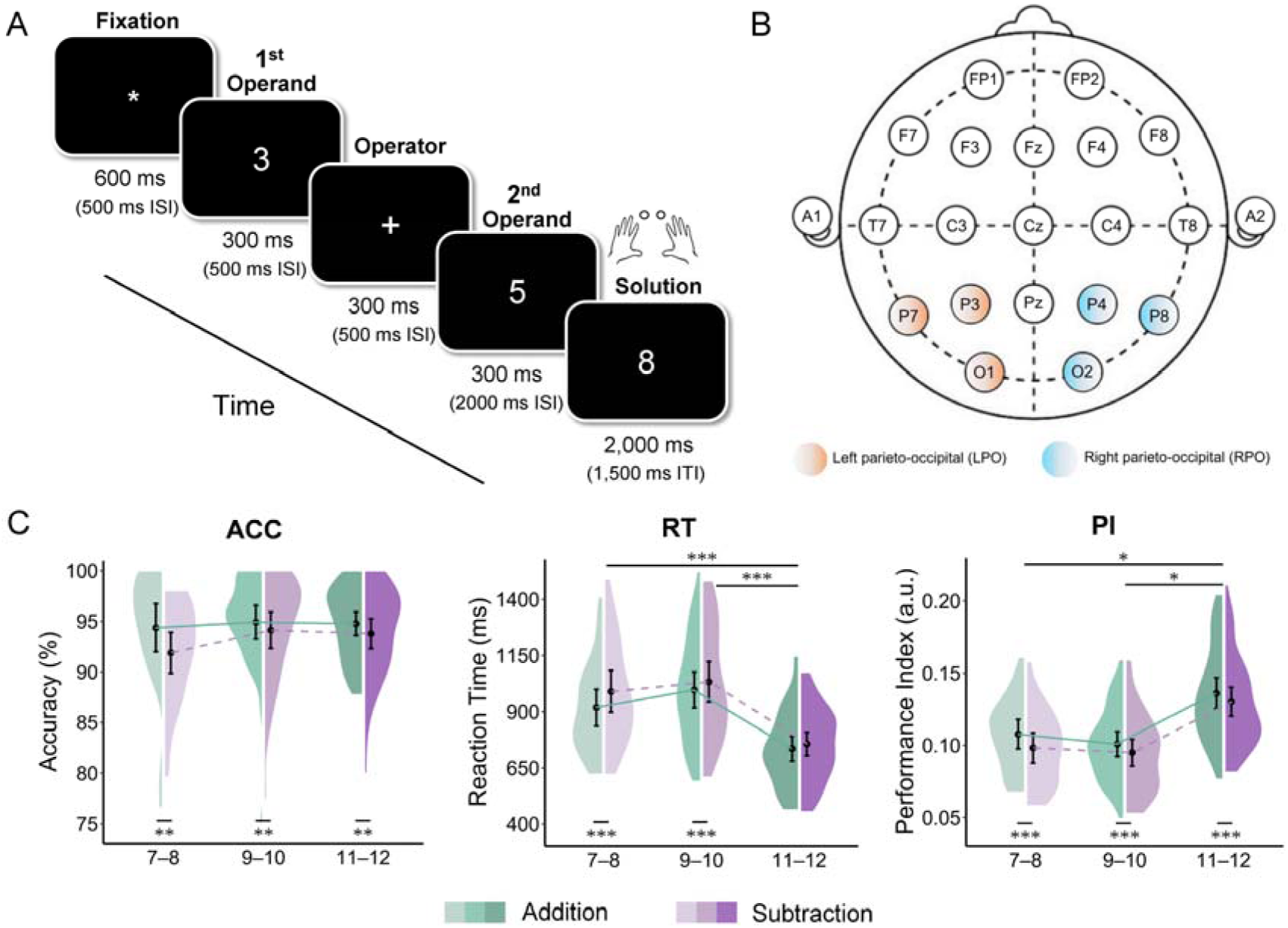
Task paradigm, sensor layout, and behavioral results. (A) Schematic representation of a single trial. All participants were instructed to justify whether the solution was congruous or incongruous with the correct answer via key presses. ISI = interstimulus interval; ITI = intertrial interval. (B) Scalp locations and electrodes used for ERP analyses. Pink and blue sections represent the left parieto-occipital (LPO: P3, P7, O1) and right parieto-occipital (RPO: P4, P8, O2) electrodes, respectively. (C) The accuracy (ACC), reaction time (RT), and performance index (PI) for the three groups (7–8-, 9–10-, and 11–12-year-olds) in the two calculations (addition [green] and subtraction [purple]). Three children with RTs exceeding ± 3 SDs from the average RT of all participants were removed from the behavioral, correlation, and mediation analyses (as below). Violin spreads present the distribution of data for all the participants. The black dots represent mean values, and the black error bars denote 95% within-subject CIs. \*\*\**p* < .001, \*\**p* < .01, \**p* < .05.

### 2.3. Behavioral analysis

The accuracy (ACC), response time (RT), and performance index (PI) were taken into behavioral analysis. For each participant, the RT was computed as the time from the solution onset until key press. Trials with incorrect or missing responses, and those with RTs exceeding ± 3 standard deviations (SDs) were excluded from the average RT (Bahnmueller et al., 2016). The ACC was calculated as the proportion of correct responses, and PI was defined as the quotient between ACC and RT (i.e., ACC/RT), with higher values indicating better arithmetic task performance (Rodriguez-Larios and Alaerts, 2019).

### 2.4. EEG recording and processing

EEG signals were acquired using a 21-channel recording system (Stellate Systems Inc., Quebec, Canada) with BESA software (MEGIS Software Co., Munich, Germany). Online EEG activity was digitized at 500 Hz. All channels were positioned on the following scalp locations according to the international 10–20 system: FP1, FP2, Fz, F3, F4, F7, F8, Cz, C3, C4, T7, T8, Pz, P3, P4, P7, P8, O1, and O2; these locations covered both the left and right earlobes (A1, A2; Fig. 1B). During data acquisition, the Cz electrode served as the online reference, and electrode impedances were kept below 5 KΩ.

The EEG recordings were preprocessed using standard processing functions from the EEGLAB toolbox (Delorme and Makeig, 2004) in MATLAB (Mathworks Inc., Natick, MA). The continuous EEG data were first downsampled to 250 Hz and re-referenced to the algebraic average of the bilateral earlobes. The data were then filtered offline between 0.5 and 40 Hz. Bad channels were replaced using a spherical spline interpolation if they exhibited artifact-laden or noisy signals for more than 40% of the entire task (Perrin et al., 1989). The unsuitable portions with large voltage drifts, head movement artifacts or paroxysmal muscle potentials were removed based on visual inspection. Independent component analysis (ICA) was then used to identify and reject components of eye movements (Delorme et al., 2001). The data were segmented into epochs from –200 to 800 ms relative to the stimulus onset (the presentation of the first or second operand). Epochs in which voltages exceeded ± 100 *µ*V in any channel were excluded. On average, 93.14 ± 7.92% of trials were retained for subsequent analyses.

### 2.5. ERP analysis

Artifact-free epochs were then time-locked to the onset of either the first or second operand, and averaged separately for each age group and calculation, using the 200-ms pe-stimulus period for baseline correction. Trials with incorrect or no responses were excluded from the ERP averaging. The P300 component was derived from the original waveforms for both operands, whereas the ΔP300 component was calculated from the difference waves obtained by subtracting the ERPs elicited by the first operand from those elicited by the second operand. For example, ΔP300 component was the difference between the P300 evoked by operand 5 (the second operand) and the P300 evoked by operand 3 (the first operand) in the trial demonstrated in Fig. 1A. Electrodes (P3, P4, P6, P8, O1, O2; Fig. 1B) and time intervals (250–350 ms) were selected using the following criteria: (1) the typical scalp location of the P300 effect reported in prior studies (Kiss et al., 1989; Takano et al., 2014; Yuan et al., 2021) and (2) visual inspection of grand-averaged waveforms and topographic maps for each age group and calculation. Both peak and mean measures were used to account for individual latency variations while reducing the sensitivity to noise. Specifically, the P300 and ΔP300 amplitudes were quantified as the mean values within a 50-ms window centered at the individual peak. For each participant, the individual peak was defined as the most positive peak value within a 100-ms time window centered at the group-averaged peak of the waveforms.

### 2.6. Statistical analyses

Statistical analyses were performed using IBM SPSS Statistics software (IBM Corp, Armonk, NY, USA). Demographic data were compared using the one-way ANOVAs for continuous values (age, IQ) and the chi-square test for categorical values (sex). One-sample *t* tests were used to evaluate whether the amplitudes of ERP components significantly differed from zero. Two-way repeated measures ANOVAs were used for behavioral indicators (ACC, RT, PI) and EEG indices (P300 amplitude, ΔP300 amplitude), with age (7–8-, 9–10-, 11–12-year-olds) as the between-subject factor, and calculation (addition, subtraction) as the within-subject factor. Greenhouse–Geisser corrections were conducted for violations of sphericity when appropriate (Geisser and Greenhouse, 1958). Significant interactions were further identified by tests of simple effects, with Bonferroni correction applied to control for the increase in type I errors. For significant comparisons, effect sizes were reported as the partial eta square (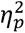) for ANOVAs and Cohen’s *d* for *t* tests.

To examine the associations between age, behavioral PI and the right parieto-occipital ΔP300 amplitude for each calculation, we used Spearman’s correlations (two-tailed) because it was less influenced by the data distributions or outliers. To further explore the role of the right parieto-occipital ΔP300 amplitude in the relationship between age and PI, we used a simple mediation model from the PROCESS macro in SPSS, with age as the independent variable, PI as the outcome variable, and the right parieto-occipital ΔP300 amplitude as the mediator. The total effect of age on PI (path *c*) is the sum of the direct effect (path *c’*) and indirect effect (path *a* × *b*) such that *c* = *c’* + (*a* × *b*). Indirect effects were assessed using bootstrap analyses (bias corrected) with 5000 samples and 95% confidence intervals (CIs). A 95% CI that does not include zero indicates a significant mediation effect (Hayes and Scharkow, 2013). Further partial correlation analyses were conducted to reveal the relationship between the right parieto-occipital ΔP300 amplitude and PI for each calculation, with age as a covariate to rule out the possible effect of age on ERPs (Böttger et al., 2002). Three children with RTs exceeding ± 3 SDs from the average RT of all participants were excluded from the behavioral, correlation, and mediation analyses. Bayes Factors (BF_01_ / BF_excl_) were calculated using JASP (JASP Team, 2025) for all the non-significant results to evaluate whether there was evidence for the null hypothesis. The significance level was set at .05.

## 3. Results

### 3.1. Behavioral results

To compare the behavioral performance of three groups of children in the two calculations, we performed the two-way repeated measures ANOVAs for ACC, RT, and PI, respectively, with group as the between-subject factor and calculation as the within-subject factor (Fig. 1C).

For ACC, the results revealed significant main effect of calculation (*F*_(1,99)_ = 8.11, *p* = .005, 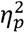 = 0.076) but no significant main effect of age (*F*_(2,99)_ = 0.92, *p* = .402, 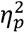 = 0.018, BF_excl_ = 5.675), suggesting that all groups of children had a similarly greater proportion of correct responses for addition (7–8-year-olds: 94.36 ± 5.86%; 9–10-year-olds: 94.92 ± 4.87%; 11–12-year-olds: 94.77 ± 3.64%) compared with subtraction (7–8-year-olds: 91.89 ± 5.02%; 9–10-year-olds: 94.13 ± 5.33%; 11–12-year-olds: 93.78 ± 4.66%). No significant interaction between group and calculation was found (*F*_(2,99)_ = 0.37, *p* = .372, 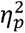 = 0.020, BF_excl_ = 1.163). For RT, a significant main effect of age was observed (*F*_(2,99)_ = 18.33, *p* < .001, 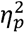 = 0.270), indicating that 11–12-year-old children showed faster responses than 7–8-year-old and 9–10-year-old children did (*p*s < .020). Furthermore, all groups of children responded more rapidly for addition (7–8-year-olds: 919 ± 204 ms; 9–10-year-olds: 997 ± 237 ms; 11–12-year-olds: 736 ± 173 ms) than for subtraction (7–8-year-olds: 991 ± 231 ms; 9–10-year-olds: 1067 ± 285 ms; 11–12-year-olds: 757 ± 164 ms), as evidenced by a significant main effect of calculation (*F*_(1,99)_ = 30.40, *p* < .001, 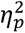 = 0.235). There was also a significant interaction between calculation and age (*F*_(2,99)_ = 3.179, *p* = .046, 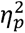 = 0.060), suggesting that 7–8- and 9–10-year-olds showed faster responses to addition than to subtraction (*p*s < .001), whereas such difference was absent in 11–12-year-olds (*p* = .181, BF_01_ = 3.770).

For PI, the results showed a significant main effect of age (*F*_(2,99)_ = 17.18, *p* < .001, 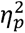 = 0.258), 11–12-year-olds performed better than 7–8- and 9–10-year-olds (*p*s < .001). Moreover, all groups of children showed better performance for addition (7–8-year-olds: 0.11 ± 0.03; 9–10-year-olds: 0.10 ± 0.03; 11–12-year-olds: 0.14 ± 0.03) compared with subtraction (7–8-year-olds: 0.10 ± 0.03; 9–10-year-olds: 0.10 ± 0.03; 11–12-year-olds: 0.13 ± 0.03), supported by a significant main effect of calculation (*F*_(1,99)_ = 33.28, *p* < .001, 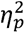 = 0.252). No significant interaction between calculation and age was found (*F*_(2,99)_ = 0.84, *p* = .433, 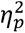 = 0.017, BF_excl_ = 1.472). The ANOVAs of all behavioral results are presented in supplementary Table A.1.

### 3.2. ERP results

3.2.1. **P300 component**

The original waveforms evoked by the second operand and the topographical maps of P300 components over the left and right parieto-occipital regions for the three age groups in the two calculations are shown in Fig. 2A and Fig. 2C. Although the early components of P1 and N1 were pronounced, statistical analysis revealed no significant difference in the amplitudes of these components between the two calculations across all age groups (*p*s > .178, BF_excl_s > 7.393).

**Fig. 2.**
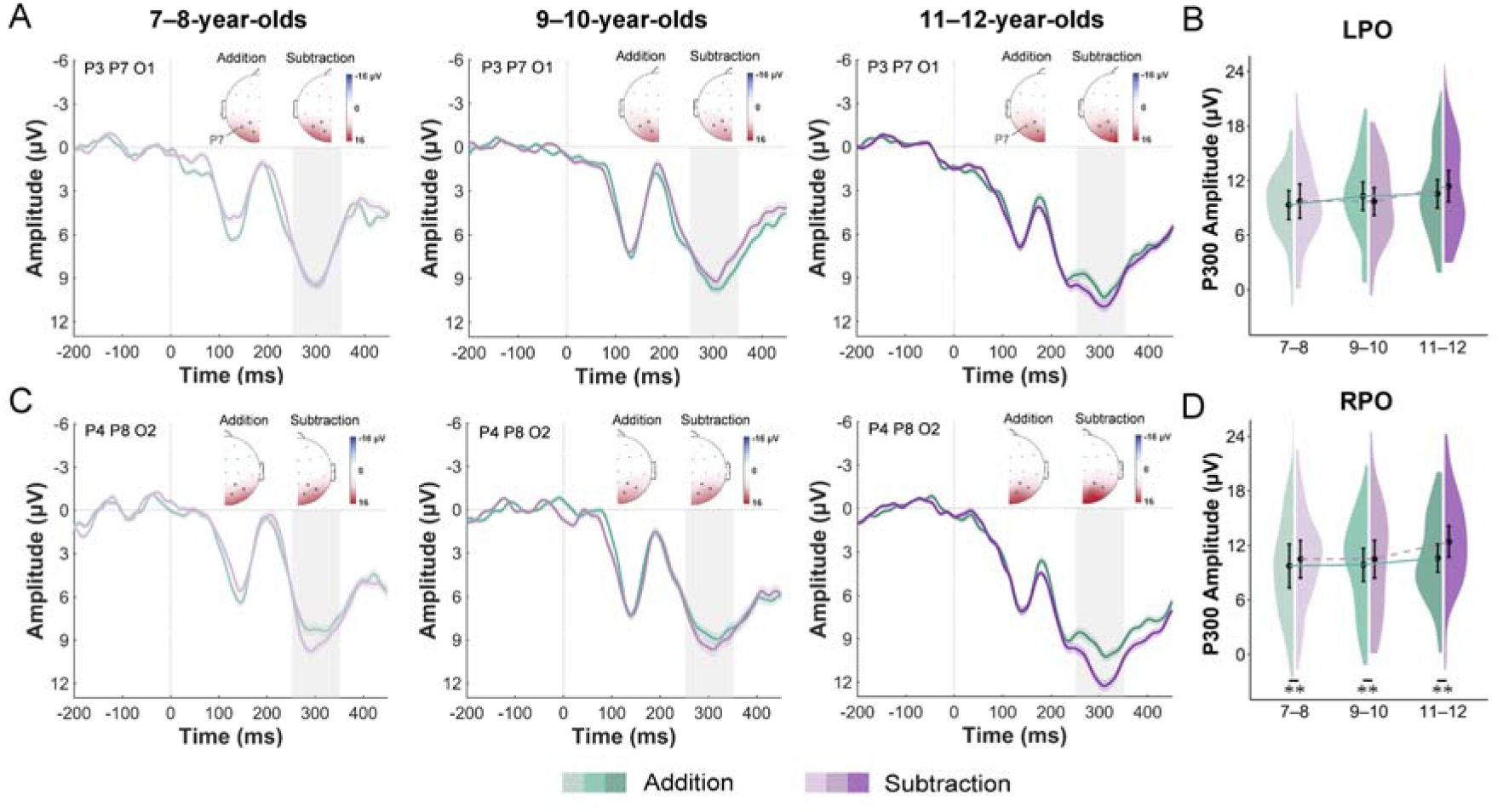
ERP results of the P300 component. (A) Grand-averaged original waveforms time-locked to the second operand and topographic maps of P300 components over the left parieto-occipital (LPO: P3, P7, O1) and (C) right parieto-occipital (RPO: P4, P8, O2) regions for the two calculations (addition [green] and subtraction [purple]) across three groups: 7–8-, 9–10-, and 11–12-year-olds. The gray shaded rectangular areas indicate the time window (250–350 ms). The black dots on the topographic maps refer to the electrodes from which the ERP waveforms were derived. (B) The P300 amplitudes between addition and subtraction in the left parieto-occipital and (D) right parieto-occipital regions across three age groups. Violin spreads present the distribution of data for all the participants. The black dots represent mean values, and the black error bars denote 95% within-subject CIs. \*\**p* < .01.

To quantify the presence of the P300 component, one-sample *t* tests were performed on the ERP waveforms for all children in each calculation, respectively. The results indicated that the mean waves of P300 (250–350 ms) were significantly elicited in both the addition (*t*_(104)_ = 20.16, *p* < .001, Cohen’s *d* = 1.968) and subtraction calculations (*t*_(104)_ = 20.18, *p* < .001, Cohen’s *d* = 1.970). Further two-way repeated measures ANOVAs were conducted separately for the left and right parieto-occipital regions to examine group differences in P300 amplitudes between the two calculations. For P300 amplitudes in the left parieto-occipital region (Fig. 2C), the main effects of calculation or age, and interaction between calculation and age were not significant (*p*s > .189, BF_excl_s > 4.224). For P300 amplitudes in the right parieto-occipital region (Fig. 2D), the results showed a significant main effect of calculation (*F*_(1,102)_ = 10.07, *p* = .002, 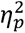 = 0.090) but no significant main effect of age (*F*_(2,102)_ = 0.82, *p* = .446,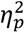 = 0.016, BF_excl_ = 0.736), indicating that all group of children exhibited greater P300 amplitudes in the right parieto-occipital region for subtraction (7–8-year-olds: 10.47 ± 5.48 μV; 9–10-year-olds: 10.48 ± 6.19 μV; 11–12-year-olds: 12.42 ± 5.44 μV) than for addition (7–8-year-olds: 9.73 ± 6.35 μV; 9–10-year-olds: 9.85 ± 5.37 μV; 11–12-year-olds: 10.58 ± 4.85 μV). No significant interaction between calculation and age was observed (*F*_(2,102)_ = 1.43, *p* = .245, 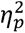 = 0.027, BF_excl_ = 5.810).

#### 3.2.2. **Δ**P300 component

The difference waves between ERPs evoked by the second operand and ERPs evoked by the first operand over the parieto-occipital regions for three groups of children in the two calculations are illustrated in Fig. 3A and Fig. 3C. The results of one-sample *t* tests against zero showed that difference waves within the 250–350 ms window were significantly more positive than zero for both addition (*t*_(104)_ = 4.31, *p* < .001, Cohen’s *d* = 0.420) and subtraction (*t*_(104)_ = 5.83, *p* < .001, Cohen’s *d* = 0.569), confirming the presence of a robust ΔP300 component.

**Fig. 3.**
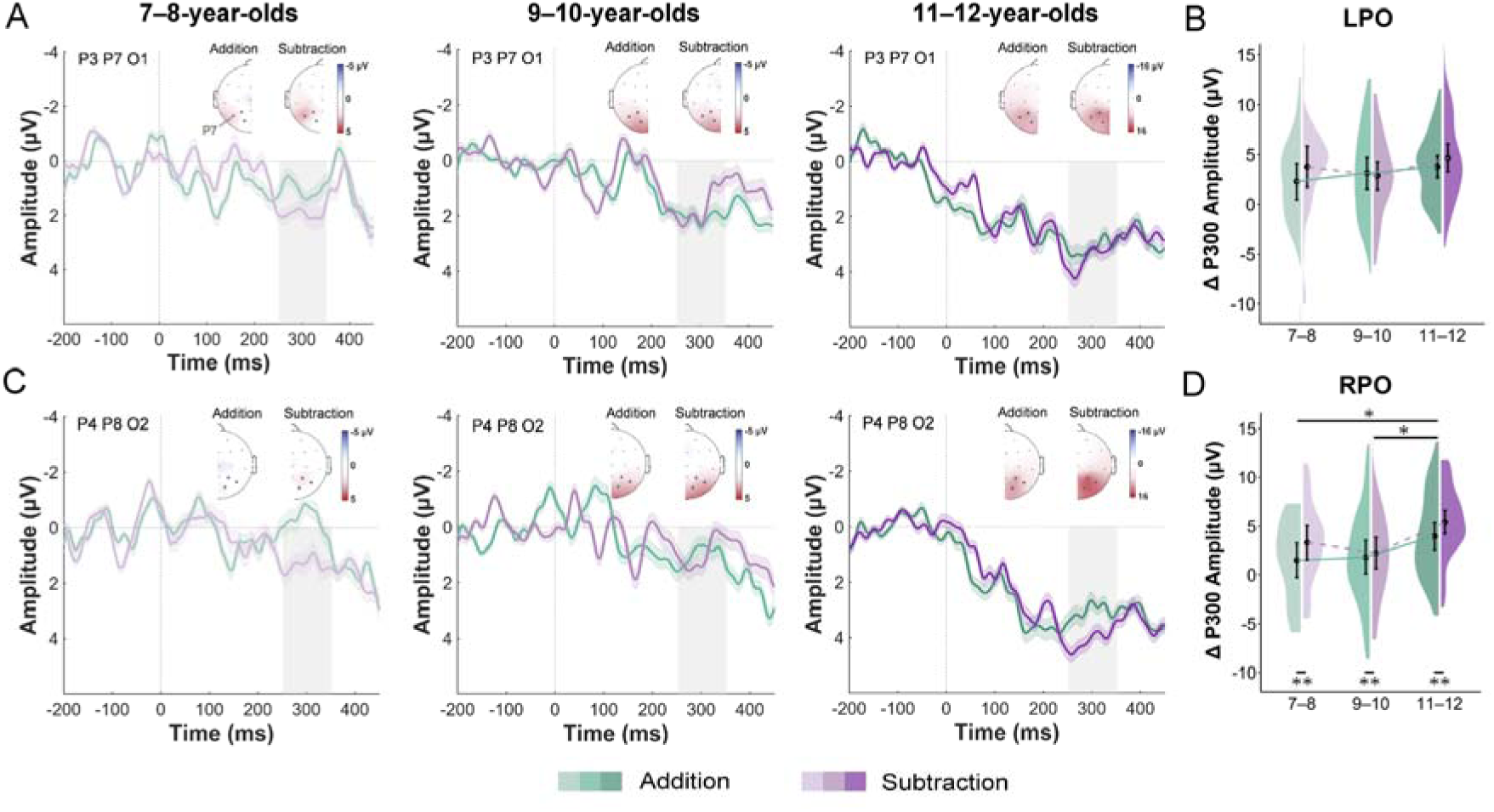
ERP results of the ΔP300 component. (A) Grand-averaged difference waveforms (ERPs evoked by the second operand minus those evoked by the first one) and topographic maps of ΔP300 components over the left parieto-occipital (LPO: P3, P7, O1) and (C) right parieto-occipital (RPO: P4, P8, O2) regions for two calculations (addition [green] and subtraction [purple]) across three groups: 7–8-, 9–10-, and 11–12-year-olds. The gray shaded rectangular areas indicate the time window (250–350 ms). The black dots on the topographic maps refer to the electrodes from which the ERP waveforms were derived. (B) The ΔP300 amplitudes between addition and subtraction in the left parieto-occipital and (D) right parieto-occipital regions across three age groups. Violin spreads present the distribution of data for all the participants. The black dots denote mean values, and the black error bars denote 95% within-subject CIs. \*\**p* < .01, \**p* < .05.

For ΔP300 amplitudes in the left parieto-occipital region (Fig. 3C), no significant main effects of calculation or age, or significant interaction between calculation and age were observed (*p*s > .190, BF_excl_s > 3.840). For ΔP300 amplitudes in the right parieto-occipital region (Fig. 3D), the results of two-way ANOVA revealed a significant main effect of age (*F*_(2,102)_ = 5.39, *p* = .006, = 0.096), suggesting that 11–12-year-olds presented significantly larger ΔP300 amplitudes than 9–10-year-olds (*p* = .012) and 7–8-year-olds (*p* = .032), similar to developmental improvements in behavioral PI. Additionally, all groups of children presented greater ΔP300 amplitudes for subtraction (7–8-year-olds: 3.41 ± 5.61 μV; 9–10-year-olds:2.24 ± 4.76 μV; 11–12-year-olds: 5.41 ± 3.73 μV) than for addition (7–8-year-olds: 0.98 ± 5.21 μV; 9–10-year-olds:1.81 ± 5.11 μV; 11–12-year-olds: 3.96 ± 4.40 μV), as indicated by a significant main effect of calculation (*F*_(1,102)_ =7.23, *p* = .008, 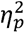 = 0.066). There was no significant interaction between calculation and age (*F*_(2,102)_ = 1.10, *p* = .338, 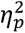 = 0.021, BF_excl_ = 1.876). The outcomes of ANOVAs for all calculations are shown in supplementary Table A.2.

### 3.3. Correlation results

To further examine whether the arithmetic-specific neural processing is related to age and behavioral performance, we performed correlation analyses using Spearman’s correlations to investigate the associations between age, PI, and right parieto-occipital ΔP300 amplitude for each calculation (Fig. 4). The results indicated that age was significantly positively correlated with PI in addition (ρ_(102)_ = 0.40, *p* < .001) and subtraction (ρ_(102)_ = 0.43, *p* < .001), suggesting age-related improvements in behavioral performance. The relationship between age and right parieto-occipital ΔP300 amplitude was significant for addition (ρ_(102)_ = 0.22, *p* = .029), and borderline significant for subtraction (ρ_(102)_ = 0.19, *p* = .059, BF_01_ = 3.322). Furthermore, right parieto-occipital ΔP300 amplitude was significantly positively correlated with PI for both addition (ρ_(102)_ = 0.24, *p* = .016) and subtraction (ρ_(102)_ = 0.20, *p* = .045), revealing that greater right parieto-occipital ΔP300 amplitudes were associated with better behavioral performance. After controlling for age, this relationship remained significant for addition (ρ_(99)_ = 0.22, *p* = .029), but was absent for subtraction (ρ_(99)_ = 0.11, *p* = .257, BF_01_ = 1.489). These findings suggest that the right parieto-occipital ΔP300 amplitude may play an important role in the age-related gains in behavioral performance, particularly for addition calculation.

**Fig. 4.**
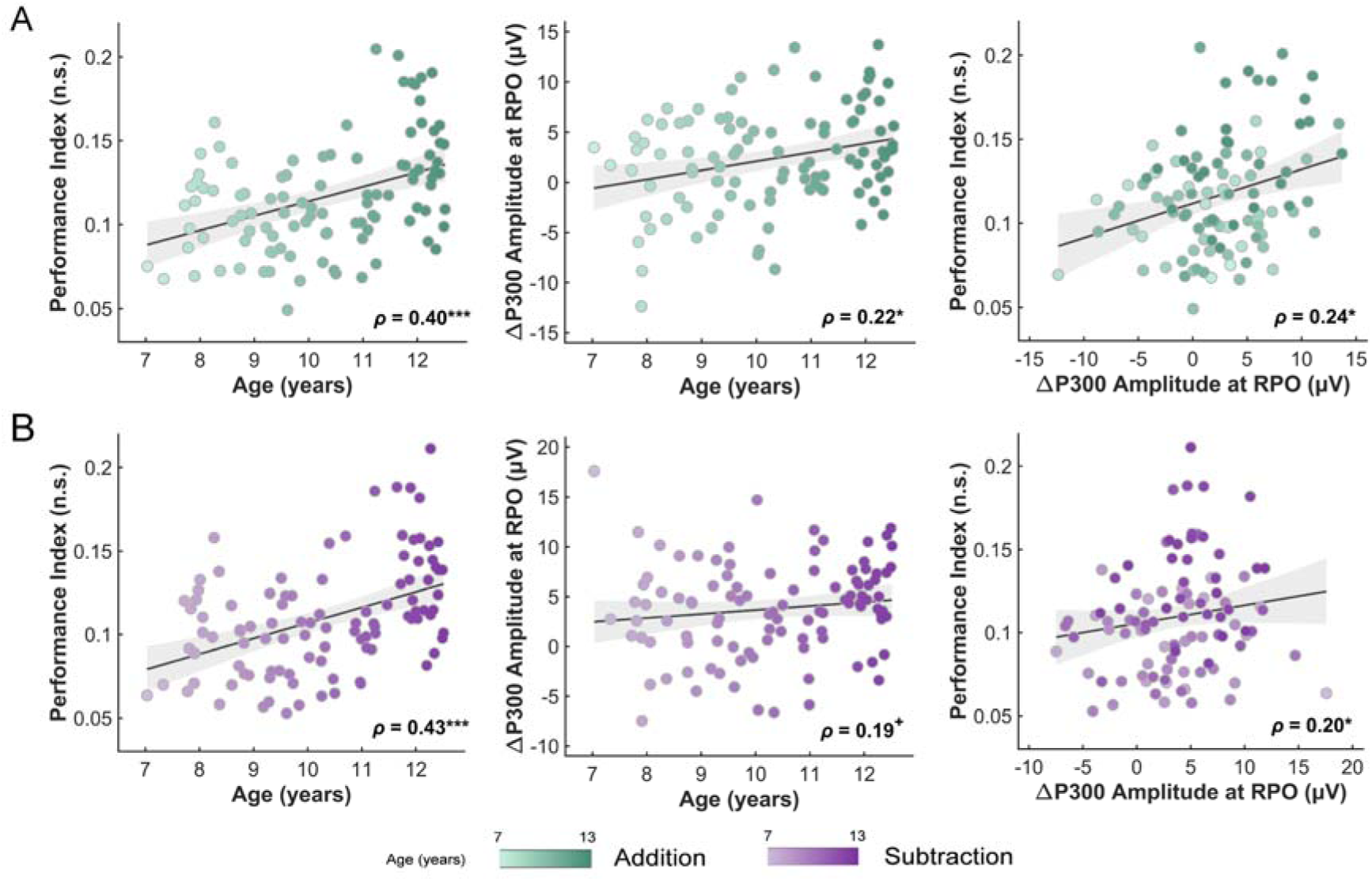
Correlation results. (A) The relationships between age, PI, and ΔP300 amplitude in the right parieto-occipital region (RPO: P4, P8, O2) for addition (green) and (B) subtraction (purple) calculations. All correlation coefficients depict Spearman’s correlations (two-tailed, uncorrected), without any factors controlled as variates. The dots refer to the individual data points. The black lines represent linear regression fits, and the gray shaded areas denote 95% CIs. \*\*\**p* < .001, \**p* < .05, ^+^ .05 ≤ *p* < .10.

### 3.4. Mediation results

Given the notable correlations observed among age, PI and right parieto-occipital ΔP300 amplitude, simple mediation model was conducted to assess whether the relationship between age and PI was mediated by right parieto-occipital ΔP300 amplitude for each calculation (Fig. 5). As shown in Fig. 5A and Fig. 5C, the results revealed a significant indirect effect of right parieto-occipital ΔP300 amplitude for addition calculation (path *c*: β = 0.009, *p* < .001, 95% CI [0.005, 0.013]; path *c’*: β = 0.008, *p* < .001, 95% CI [0.004, 0.011]; path *a* × *b*: β = 0.001, 95% CI [0.001, 0.003]), suggesting that right parieto-occipital ΔP300 amplitude partially mediated the positive relation between age and PI (path *a*: β = 0.266, *p* = .007, 95% CI [0.238, 1.466]; path *b*: β = 0.205, *p* = .029, 95% CI [0.001, 0.003]). In contrast, no significant indirect effect of right parieto-occipital ΔP300 amplitude between age and PI was observed for subtraction calculation (95% CIs included zero; Fig. 5B and Fig. 5D). The outputs for all mediation models are listed in supplementary Table A.3.

**Fig. 5.**
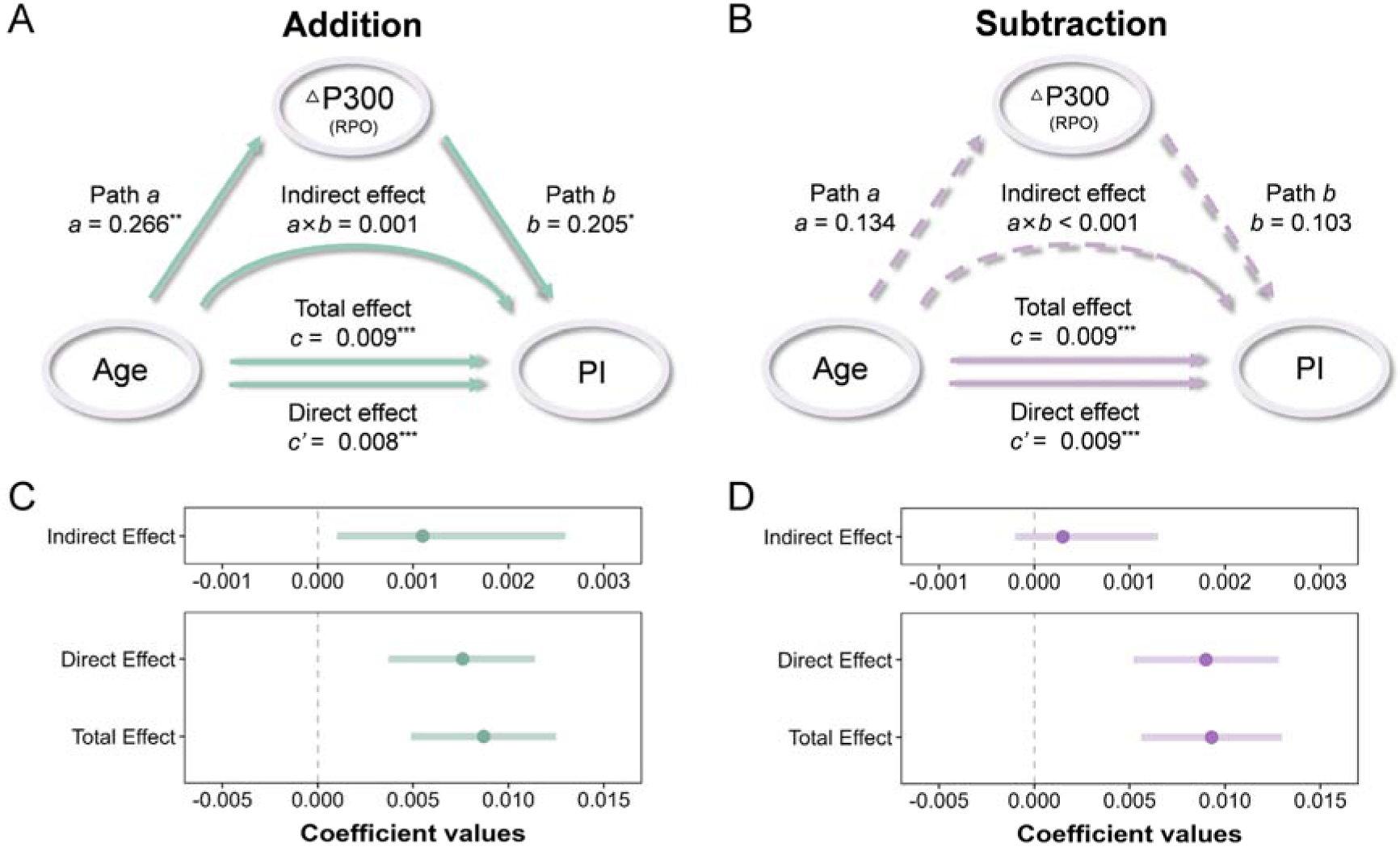
Mediation results. (A) Mediation models of the right parieto-occipital (RPO: P4, P8, O2) ΔP300 amplitude between age and PI for addition (green) and (B) subtraction (purple) calculations. Paths *a*, *b*, and *c* represent standardized beta coefficients of the direct path strength, and path *c’* denotes the standardized beta coefficient of path strength after controlling for the right parieto-occipital ΔP300 amplitude. The solid gray lines indicate significant effects and the dashed gray lines denote non-significant effects. RPO = right parieto-occipital (P4, P8, O2). (C) The effect estimates and 95% CIs for addition and (D) subtraction calculations. The red dots indicate standardized beta coefficients, the orange solid lines represent 95% CIs, and the blue dashed lines denote zero. \*\*\**p* < .001, \*\**p* < .01, \**p* < .05.

## 4. Discussion

Using an arithmetic task with item-by-item presentation, we investigate developmental trajectories of addition and subtraction calculations in children aged 7–12 years. By applying a difference wave method (second minus first operand), we were able to isolate arithmetic-specific brain activities (as indexed by the ΔP300 amplitude) from general cognitive processes related to two operands. Behaviorally, children calculated addition faster and more accurately than subtraction; ERP results revealed that children exhibited greater right parieto-occipital P300 and ΔP300 amplitudes in subtraction than in addition. Specifically, 11–12-year-olds showed better behavioral performance and greater right parieto-occipital ΔP300 amplitudes compared with 7–8- and 9–10-year-olds. The greater right parieto-occipital ΔP300 amplitude predicted better behavioral performance in both addition and subtraction, whereas it mediated the relationship between age and behavioral performance only in addition. These findings provide novel and neurophysiological evidence for the development of addition and subtraction during childhood and highlight the right parieto-occipital ΔP300 as a potential neural marker of arithmetic performance, especially for addition calculation.

All children solved subtraction problems slower and made more mistakes than addition problems, possibly because addition is more commonly and frequently used in our daily life, whereas subtraction is relatively less automated to grasp (Campbell, 2005). Moreover, older children calculated addition and subtraction problems more efficiently than younger children. Although we did not find a significant age-related increase in ACC, which aligns with previous findings reporting stable accuracy but faster RT in older children during two-operand arithmetic calculations (Rivera et al., 2005), this may be due to a ceiling effect. The better behavioral performance in older children may be result from the neural maturation that facilitates the developmental shift in strategy use: with increasing age, children are more likely to use retrieval or decomposition strategies and less likely to use procedural strategies such as counting (Torbeyns et al., 2004). That is, younger children may count ‘‘4, 5, 6’’ to solve the problem “4 +2”. As children grow up, they tend to form a long-term memory association between the answer (‘‘6’) and problem (‘‘4 + 2’’), thereby retrieving the solution directly from memory and calculating faster (Caviola et al., 2018; Siegler and Shrager, 1984). Notably, we observed that 11–12 years performed both calculations better than younger children, indicating a key developmental age of 11–12 years for neural and behavioral maturation of strategy use.

We found that all groups of children exhibited greater P300 amplitudes in subtraction than addition. Arithmetic has been shown to depend on widespread neural areas involved in overall cognitive processes, including general (e.g., language, working memory, long term memory, and visuo-spatial abilities) and domain-specific processes (Arsalidou et al., 2018). When the second operand appears, children begin to perform the arithmetic calculation; thus, the P300 evoked by the second operand may be a neural marker of overall cognitive processes engaged in arithmetic (Gao et al., 2022; Jost et al., 2004). As such, the larger P300 amplitudes in subtraction may reflect a greater level of overall cognitive resources required when solving subtraction problems compared with addition. This P300 difference between the two calculations was manifested in the right parieto-occipital region, aligning with previous studies showing stronger activation for subtraction than addition in the right posterior parietal cortex (Kawashima et al., 2004; Rosenberg-Lee et al., 2011). It is possible that subtraction relies more overall resources from the right-lateralized region for the abstract representation and procedural manipulation of numerical magnitude (Holloway and Ansari, 2010; Prado et al., 2014).

The most novel and critical finding of our study is the ΔP300 amplitude, which is thought to reflect arithmetic-specific processes after subtracting general processes elicited by the two operands. In line with the age-related improvements in behavioral PI, we found that older children presented greater right parieto-occipital ΔP300 amplitudes, indicating that the arithmetic-specific neural processing matures with age (Dehaene et al., 2003). Our results suggest that this neural maturation has occurred from an early age of 7–8 years, which is earlier than previously assumed (Dong et al., 2007), reaching a relatively stable level through 11–12 years. Compared with subtraction, all children exhibited smaller right parieto-occipital ΔP300 amplitudes in addition, which played a mediation role in the relationship between age and PI. This pattern may be explained by the mental-attentional theory (Pascual-Leone and Johnson, 2011), positing that the right hemisphere could be mobilized in search of potentially useful overlearned or automatized schemes when mental attentional demand exceeds the processing capacity of the left hemisphere. Accordingly, compared with younger children, older children are better able to recruit additional right-hemisphere resources for spatial-numerical processes, resulting in improved behavioral performance in addition calculation. These findings suggest distinct roles of the right parieto-occipital ΔP300 in addition and subtraction calculations: addition shows an age-related increase in the mobilization of right parieto-occipital cognitive resources for arithmetic-specific processes, whereas subtraction is so demanding that children continue to engage a similarly high level of both general and arithmetic-specific cognitive processes even as they grow older.

There are still some limitations in the current study. First, we used an arithmetic task involved only calculations using Arabic numerals with a limited range from 2 to 20. It may not fully capture the complexity and diversity of arithmetic processing in natural educational settings. Future studies are encouraged to use real-world math tasks involved larger numbers, multi-step reasoning, or word problems that engage broader arithmetic cognitive processes. Secondly, children who learn ideographic systems (e.g., Chinese) may develop different neural pathways relative to children who use alphabetic systems. Future neuroimaging studies are necessary to compare these groups and provide deeper insight into the cultural specificity of arithmetic-specific neural mechanisms. Finally, we used the cross-sectional design that limited our interpretations on uncertain developmental changes at the individual level. It seems promising for future longitudinal research to track additional influencing factors and to assess the stability of neural markers over time within individuals.

## 5. Conclusion

This study investigates distinct developmental trajectories of addition and subtraction in school-aged children, highlighting the right parieto-occipital ΔP300 as a neural marker of arithmetic performance, particularly for addition calculation. The results revealed that 11–12 years was a key developmental stage for both behavioral and neural maturation in arithmetic. Subtraction required consistently more general and arithmetic-specific resources from the right parieto-occipital region over development, whereas addition showed age-related increase of arithmetic-specific resources that facilitates behavioral performance. These findings provide neurophysiology evidence for the maturation of arithmetic calculation during childhood.

## Data Availability

The data, analytical codes, and materials are available on the request.

## CRediT Authorship Contribution Statement

**Manqi Zhou**: Conceptualization, Investigation, Methodology, Formal analysis, Visualization, Writing – original draft, Writing – review & editing.

**Shiyan Ji**: Date curation, Investigation, Validation, Software.

**Yalun Zhang**: Investigation, Writing – original draft, Writing – review & editing.

**Huijuan Shen**: Date curation, Investigation. **Jipeng Huang:** Supervision, Investigation. **Qinfen Zhang**: Date curation, Validation.

**Yiwen Li**: Supervision, Writing – review & editing.

**Haitian Mei**: Date curation. **Rong Kuang**: Date curation. **Yuanxin Lin**: Date curation.

**Yan Song**: Supervision, Funding acquisition, Writing – review & editing.

**Jing Huang**: Supervision, Funding acquisition, Writing – review & editing.

**Xuan Dong**: Resources, Supervision, Project administration, Funding acquisition.

## Acknowledgements

This work was supported by the STI 2030—Major Projects [No.2021ZD0200500 to Y.S.]; the National Natural Science Foundation of China [No. 32271094 to Y.S.; No. 32200870 to J.H.]; the National Key Research and Development Program of China [No. 2024YFF0509100 to J.H.]; and the Nantong University Special Research Fund for Clinical Medicine [No. 2024YL033 to X.D.; No. 2024LQ035 to X.D]. No generative AI and AI-assisted technologies were used in the manuscript preparation process.

## Ethics statement

This work was approved by the Ethics Committee of Children’s Hospital of Changzhou according to the Declaration of Helsinki. Informed consent was obtained from all participants involved in the study.

## References

Anderson, J., 2000. Cognitive psychology and its implications, 5th ed. ed. New York: Worth Publishers, New York, NY.

Arsalidou, M., Pawliw-Levac, M., Sadeghi, M., Pascual-Leone, J., 2018. Brain areas associated with numbers and calculations in children: meta-analyses of fMRI studies. Dev. Cogn. Neurosci. 30, 239–250. 10.1016/j.dcn.2017.08.002.

Bahnmueller, J., Huber, S., Nuerk, H.-C., Göbel, S.M., Moeller, K., 2016. Processing multi-digit numbers: a translingual eye-tracking study. Psychol. Res. 80, 422–433. 10.1007/s00426-015-0729-y.

Blankenberger, S., 2001. The arithmetic tie effect is mainly encoding-based. Cognition 82, B15–B24. 10.1016/S0010-0277(01)00140-8.

Böttger, D., Herrmann, C.S., von Cramon, D.Y., 2002. Amplitude differences of evoked alpha and gamma oscillations in two different age groups. Int. J. Psychophysiol. 45, 245–251. 10.1016/S0167-8760(02)00031-4.

Butterworth, B., 2005. The development of arithmetical abilities. J. Child Psychol. Psychiatry 46, 3–18. 10.1111/j.1469-7610.2004.00374.x.

Butterworth, B., Varma, S., Laurillard, D., 2011. Dyscalculia: from brain to education. Science 332, 1049–1053. 10.1126/science.1201536.

Campbell, J.I.D., 2005. The handbook of mathematical cognition. New York: Psychology Press, New York. 10.4324/9780203998045.

Campbell, J.I.D., Graham, D.J., 1985. Mental multiplication skill: Structure, process, and acquisition. Can. J. Psychol. Rev. Can. Psychol. 39, 338–366. 10.1037/h0080065.

Caviola, S., Mammarella, I.C., Pastore, M., LeFevre, J.-A., 2018. Children’s strategy choices on complex subtraction problems: individual differences and developmental changes. Front. Psychol. 9. 10.3389/fpsyg.2018.01209.

Chassy, P., Grodd, W., 2012. Comparison of quantities: core and format-dependent regions as revealed by fMRI. Cereb. Cortex 22, 1420–1430. 10.1093/cercor/bhr219.

Chochon, F., Cohen, L., Moortele, P.F. van de Dehaene, S., 1999. Differential contributions of the left and right inferior parietal lobules to number processing. J. Cogn. Neurosci. 11, 617–630. 10.1162/089892999563689.

De Smedt, B., Grabner, R.H., Studer, B., 2009. Oscillatory EEG correlates of arithmetic strategy use in addition and subtraction. Exp. Brain Res. 195, 635–642. 10.1007/s00221-009-1839-9.

Dehaene, S., Cohen, L., 1997. Cerebral pathways for calculation: double dissociation between rote verbal and quantitative knowledge of arithmetic. Cortex 33, 219–250. 10.1016/S0010-9452(08)70002-9.

Dehaene, S., Piazza, M., Pinel, P., Cohen, L., 2003. Three parietal circuits for number processing. Cogn. Neuropsychol. 20, 487–506. 10.1080/02643290244000239.

Delorme, A., Makeig, S., 2004. EEGLAB: an open source toolbox for analysis of single-trial EEG dynamics including independent component analysis. J. Neurosci. Methods 134, 9–21. 10.1016/j.jneumeth.2003.10.009.

Delorme, A., Makeig, S., Sejnowski, T., 2001. Automatic artifact rejection for EEG data using high-order statistics and independent component analysis. Paper Presented at the Proc. of the 3rd International Workshop on ICA, 457–462.

Dong, X., Wang, S., Yang, Yilin, Ren, Y., Meng, P., Yang, Yuxia, 2007. Cognitive development of semantic process and mental arithmetic in childhood: an event-related potential. Data Sci. J. 6, S535–S547. 10.2481/dsj.6.S535.

Evans, T.M., Flowers, D.L., Luetje, M.M., Napoliello, E., Eden, G.F., 2016. Functional neuroanatomy of arithmetic and word reading and its relationship to age. NeuroImage 143, 304–315. 10.1016/j.neuroimage.2016.08.048.

Faul, F., Erdfelder, E., Lang, A.-G., Buchner, A., 2007. G*power 3: a flexible statistical power analysis program for the social, behavioral, and biomedical sciences. Behav. Res. Methods 39, 175–191. 10.3758/BF03193146.

Formoso, J., Injoque-Ricle, I., Barreyro, J.-P., Calero, A., Jacubovich, S., Burín, D.I., 2018. Mathematical cognition, working memory, and processing speed in children. Cogn. Brain Behav. Interdiscip. J. 22, 59–84. 10.24193/cbb.2018.22.05.

Gao, Y., Wang, X., Huang, B., Li, H., Wang, Y., Si, J., 2022. How numerical surface forms affect strategy execution in subtraction? Evidence from behavioral and ERP measures. Exp. Brain Res. 240, 439–451. 10.1007/s00221-021-06259-6.

Geisser, S., Greenhouse, S.W., 1958. An extension of box’s results on the use of the F distribution in multivariate analysis. Ann. Math. Stat. 29, 885–891. 10.1214/aoms/1177706545.

Hayes, A.F., Scharkow, M., 2013. The relative trustworthiness of inferential tests of the indirect effect in statistical mediation analysis: does method really matter? Psychol. Sci. 24, 1918–1927. 10.1177/0956797613480187.

Holloway, I.D., Ansari, D., 2010. Developmental specialization in the right intraparietal sulcus for the abstract representation of numerical magnitude. J. Cogn. Neurosci. 22, 2627–2637. 10.1162/jocn.2009.21399.

Ischebeck, A., Zamarian, L., Schocke, M., Delazer, M., 2009. Flexible transfer of knowledge in mental arithmetic — an fMRI study. NeuroImage 44, 1103–1112. 10.1016/j.neuroimage.2008.10.025.

Jasinski, E.C., Coch, D., 2012. ERPs across arithmetic operations in a delayed answer verification task. Psychophysiology 49, 943–958. 10.1111/j.1469-8986.2012.01378.x.

JASP Team, 2025. JASP (Version 0.17.1), [Computer Software]. The Netherlands: University of Amsterdam. https://jasp-stats.org/.

Jost, K., Beinhoff, U., Hennighausen, E., Rösler, F., 2004. Facts, rules, and strategies in single-digit multiplication: evidence from event-related brain potentials. Cogn. Brain Res. 20, 183–193. 10.1016/j.cogbrainres.2004.02.005.

Kaufmann, L., Wood, G., Rubinsten, O., Henik, A., 2011. Meta-analyses of developmental fMRI studies investigating typical and atypical trajectories of number processing and calculation. Dev. Neuropsychol. 36, 763–787. 10.1080/87565641.2010.549884.

Kawashima, R., Taira, M., Okita, K., Inoue, K., Tajima, N., Yoshida, H., Sasaki, T., Sugiura, M., Watanabe, J., Fukuda, H., 2004. A functional MRI study of simple arithmetic—a comparison between children and adults. Cogn. Brain Res. 18, 227–233. 10.1016/j.cogbrainres.2003.10.009.

Kiss, I., Dashieff, R.M., Lordeon, P., 1989. A parietooccipital generator for P300: evidence from human intracranial recordings. Int. J. Neurosci. 49, 133–139. 10.3109/00207458909087048.

Kong, J., Wang, C., Kwong, K., Vangel, M., Chua, E., Gollub, R., 2005. The neural substrate of arithmetic operations and procedure complexity. Cogn. Brain Res. 22, 397–405. 10.1016/j.cogbrainres.2004.09.011.

LeFevre, J.-A., Liu, J., 1997. The role of experience in numerical skill: multiplication performance in adults from Canada and China. Math. Cogn. 3, 31–62. 10.1080/135467997387470.

Menon, V., 2010. Developmental cognitive neuroscience of arithmetic: implications for learning and education. ZDM 42, 515–525. 10.1007/s11858-010-0242-0.

Montefinese, M., Turco, C., Piccione, F., Semenza, C., 2017. Causal role of the posterior parietal cortex for two-digit mental subtraction and addition: a repetitive TMS study. NeuroImage 155, 72–81. 10.1016/j.neuroimage.2017.04.058.

Muluh, E.T., Vaughan, C.L., John, L.R., 2011. High resolution event-related potentials analysis of the arithmetic-operation effect in mental arithmetic. Clin. Neurophysiol. 122, 518–529. 10.1016/j.clinph.2010.08.008.

Núñez-Peña, M.I., Honrubia-Serrano, M.L., Escera, C., 2004. Problem size effect in additions and subtractions: an event-related potential study. Neurosci. Lett. 373, 21–25. 10.1016/j.neulet.2004.09.053.

Pascual-Leone, J., Johnson, J., 2011. A developmental theory of mental attention: its application to measurement and task analysis, in: Cognitive Development and Working Memory: A Dialogue between Neo-Piagetian Theories and Cognitive Approaches. Psychology Press, New York, NY, US, pp. 13–46.

Perrin, F., Pernier, J., Bertrand, O., Echallier, J.F., 1989. Spherical splines for scalp potential and current density mapping. Electroencephalogr. Clin. Neurophysiol. 72, 184–187. 10.1016/0013-4694(89)90180-6.

Polich, J., 2007. Updating P300: an integrative theory of P3a and P3b. Clin. Neurophysiol. 118, 2128–2148. 10.1016/j.clinph.2007.04.019.

Prado, J., Mutreja, R., Booth, J.R., 2014. Developmental dissociation in the neural responses to simple multiplication and subtraction problems. Dev. Sci. 17, 537–552. 10.1111/desc.12140.

Prado, J., Mutreja, R., Zhang, H., Mehta, R., Desroches, A.S., Minas, J.E., Booth, J.R., 2011. Distinct representations of subtraction and multiplication in the neural systems for numerosity and language. Hum. Brain Mapp. 32, 1932–1947. 10.1002/hbm.21159.

Price, G.R., Holloway, I., Räsänen, P., Vesterinen, M., Ansari, D., 2007. Impaired parietal magnitude processing in developmental dyscalculia. Curr. Biol. 17, R1042–R1043. 10.1016/j.cub.2007.10.013.

Rivera, S.M., Reiss, A.L., Eckert, M.A., Menon, V., 2005. Developmental changes in mental arithmetic: evidence for increased functional specialization in the left inferior parietal cortex. Cereb. Cortex 15, 1779–1790. 10.1093/cercor/bhi055.

Rodriguez-Larios, J., Alaerts, K., 2019. Tracking transient changes in the neural frequency architecture: harmonic relationships between theta and alpha peaks facilitate cognitive performance. J. Neurosci. 39, 6291–6298. 10.1523/JNEUROSCI.2919-18.2019.

Rosenberg-Lee, M., Chang, T.T., Young, C.B., Wu, S., Menon, V., 2011. Functional dissociations between four basic arithmetic operations in the human posterior parietal cortex: a cytoarchitectonic mapping study. Neuropsychologia 49, 2592–2608. 10.1016/j.neuropsychologia.2011.04.035.

Salillas, E., Benavides-Varela, S., Semenza, C., 2023. The brain lateralization and development of math functions: progress since sperry, 1974. Front. Hum. Neurosci. 17, 1288154. 10.3389/fnhum.2023.1288154.

Siegler, R., Shrager, J., 1984. Strategy choices in addition and subtractionL: how do children know what to do?, in: Origins of Cognitive Skills, The Eighteenth Annual Carnegie Symposlum on Cognition. London: Lawrence Erlbaum Associates, London, pp. 229–293.

Takano, K., Ora, H., Sekihara, K., Iwaki, S., Kansaku, K., 2014. Coherent activity in bilateral parieto-occipital cortices during P300-BCI operation. Front. Neurol. 5, 74. 10.3389/fneur.2014.00074.

Torbeyns, J., Verschaffel, L., Ghesquière, P., 2004. Strategic aspects of simple addition and subtraction: the influence of mathematical ability. Learn. Instr. 14, 177–195. 10.1016/j.learninstruc.2004.01.003.

Wechsler, D., 1974. Manual for the wechsler intelligence scale for children (revised). New York: Psychological Corporation, New York.

Yuan, G., Liu, G., Wei, D., 2021. Roles of P300 and late positive potential in initial romantic attraction. Front. Neurosci. 15, 718847. 10.3389/fnins.2021.718847.

Zhou, X., Chen, Chuansheng, Dong, Q., Zhang, H., Zhou, R., Zhao, H., Chen, Chunhui, Qiao, S., Jiang, T., Guo, Y., 2006. Event-related potentials of single-digit addition, subtraction, and multiplication. Neuropsychologia 44, 2500–2507. 10.1016/j.neuropsychologia.2006.04.003.

