## Supplementary material for "Developmental Trajectories of Addition and Subtraction in Early School-aged Children": Table A.1; Table A.2; Table A.3

**RH** = Development of Calculation in Children

***Supplementary Materials***

**Table A.1**

Inferential statistics derived from mixed factorial ANOVAs for behavior indicators (ACC, RT, and PI).

|  | A. 7–8-year-olds | | B. 9–10-year-olds | | C. 11–12-year-olds | | Group  (A vs. B vs. C) | Calculation  (① vs. ②) |  | *F* | *p-*value | $\eta_{p}^{2}$ |
| --- | --- | --- | --- | --- | --- | --- | --- | --- | --- | --- | --- | --- |
|  | ① Add^a^ | ② Sub^b^ | ① Add^a^ | ② Sub^b^ | ① Add^a^ | ② Sub^b^ |  |  |  |  |  |  |
| ACC | 94.37 | 91.89 | 94.92 | 94.13 | 94.77 | 93.78 | n.s. | **①>②^**^** | Calculation | 8.11 | **.005^**^** | 0.076 |
|  | (5.87) | (5.02) | (4.87) | (5.33) | (3.64) | (4.66) |  |  | Group | 0.92 | .402 | 0.018 |
|  | n.s. | | n.s. | | n.s. | |  |  | Group × Calculation | 1.00 | .372 | 0.020 |
| RT | 918.62 | 990.67 | 997.38 | 1067.04 | 735.79 | 756.60 | **A>C^***^** | **①<②^***^** | Calculation | 30.40 | **<.001^***^** | 0.235 |
|  | (203.95) | (230.67) | (236.76) | (285.46) | (173.09) | (164.08) | **B>C^***^** |  | Group | 18.33 | **<.001^***^** | 0.270 |
|  | **①<②^***^** | | **①<②^***^** | | n.s. | |  |  | Group × Calculation | 3.18 | **.046^*^** | 0.060 |
| PI | 0.11 | 0.10 | 0.10 | 0.09 | 0.14 | 0.13 | **A<C^***^** | **①>②^***^** | Calculation | 33.28 | **<.001^***^** | 0.252 |
|  | (0.03) | (0.03) | (0.03) | (0.03) | (0.03) | (0.03) | **B<C^***^** |  | Group | 17.18 | **<.001^***^** | 0.258 |
|  | n.s. | | n.s. | | n.s. | |  |  | Group × Calculation | 0.84 | .433 | 0.017 |

*Note*. For each behavioral measure, the means are presented in the first row, and the standard deviations (SDs) are given in the second row. The third row reports the results of simple effect analyses for Calculations (addition, subtraction) within Groups (7–8-, 9–10-, 11–12-year-olds) when interactions between Group and Calculation are significant; otherwise, they are marked as n.s. (not significant). The right half of the table presents the significance of main effects of Group and Calculation through post hoc tests. The values in bold indicate significant effects after applying a Bonferroni correction. ACC = accuracy; RT = mean reaction time for correct trials; PI = performance index (ACC/RT).

^a^ represents the addition calculation. ^b^ represents the subtraction calculation. ^***^*p* < .001, ^**^*p* < .01, ^*^*p* < .05.

**Table A.2**

Inferential statistics derived from mixed ANOVAs for EEG indices (P300 amplitude from original waves, and ∆P300 amplitude from difference waves).

|  | A. 7–8-year-olds | | B. 9–10-year-olds | | C. 11–12-year-olds | | Group  (A vs. B vs. C) | Calculation  (① vs. ②) |  | *F* | *p-*value | $\eta_{p}^{2}$ |
| --- | --- | --- | --- | --- | --- | --- | --- | --- | --- | --- | --- | --- |
|  | ① Add^a^ | ② Sub^b^ | ① Add^a^ | ② Sub^b^ | ① Add^a^ | ② Sub^b^ |  |  |  |  |  |  |
| P300^a^ | 9.92 | 9.73 | 10.27 | 9.68 | 10.53 | 11.37 | n.s. | n.s. | Calculation | 0.01 | .949 | 0.000 |
|  | (5.32) | (4.99) | (4.66) | (4.53) | (4.90) | (5.39) |  |  | Group | 0.63 | .534 | 0.012 |
|  | n.s. | | n.s. | | n.s. | |  |  | Group × Calculation | 1.69 | .189 | 0.032 |
| P300^b^ | 9.73 | 10.47 | 9.85 | 10.48 | 10.58 | 12.42 | n.s. | **①<②^**^** | Calculation | 10.07 | **.002^**^** | 0.090 |
|  | (6.35) | (5.48) | (5.37) | (6.19) | (4.85) | (5.44) |  |  | Group | 0.82 | .446 | 0.016 |
|  | n.s. | | n.s. | | n.s. | |  |  | Group × Calculation | 1.43 | .245 | 0.027 |
| ∆P300^a^ | 2.28 | 3.76 | 3.12 | 2.86 | 4.17 | 4.66 | n.s. | n.s. | Calculation | 1.28 | .261 | 0.012 |
|  | (4.77) | (5.43) | (4.71) | (4.16) | (4.27) | (4.39) |  |  | Group | 1.69 | .190 | 0.032 |
|  | n.s. | | n.s. | | n.s. | |  |  | Group × Calculation | 0.94 | .394 | 0.018 |
| ∆P300^b^ | 0.98 | 3.41 | 1.81 | 2.24 | 3.96 | 5.41 | **A<C^*^** | **①<②^**^** | Calculation | 7.23 | **.008^**^** | 0.066 |
|  | (5.21) | (5.61) | (5.11) | (4.76) | (4.40) | (3.73) | **B<C^*^** |  | Group | 5.39 | **.006^**^** | 0.096 |
|  | n.s. | | n.s. | | n.s. | |  |  | Group × Calculation | 1.10 | .338 | 0.021 |

*Note.* For each EEG indicator, the means are presented in the first row, and the standard deviations (SDs) are given in the second row. The third row reports the results of simple effect analyses for Calculations (addition, subtraction) within Groups (7–8-, 9–10-, 11–12-year-olds) when interactions between Group and Calculation are significant; otherwise, they are marked as n.s. (not significant). The right half of the table presents the significance of main effects of Group and Calculation through post hoc tests. The values in bold indicate significant effects after a Bonferroni correction.

^a^ represents the left parieto-occipital region (P3, P7, O1). ^b^ represents the right parieto-occipital region (P4, P8, O2). ^**^*p* < .01, ^*^*p* < .05.

**Table A.3**

Estimated path coefficients and significance tests of the simple mediation analyses (model 4) assessing the direct and indirect effects for age on PI, with the ∆P300 amplitude in the right parieto-occipital region as the mediator variable.

|  |  |  |  |  | 95% confidence interval (CI) | |  |  |
| --- | --- | --- | --- | --- | --- | --- | --- | --- |
|  | Type | Effect | β | SE | Lower | Upper | *t* | *p-*value |
| **Addition** | Indirect | Age → ∆P300 amplitude → PI (path *a × b*) | 0.001 | 0.001 | 0.001 | 0.003 | - | - |
|  | Component | Age → ∆P300 amplitude (path *a*) | 0.266 | 0.309 | 0.238 | 1.466 | 2.753 | **.007^**^** |
|  |  | ∆P300 amplitude → PI (path *b*) | 0.205 | 0.001 | 0.001 | 0.003 | 2.217 | **.029^*^** |
|  | Direct | Age → PI (path *c’*) | 0.007 | 14.62 | 0.004 | 0.011 | 3.903 | **<.001^***^** |
|  | Total | Age → PI (path *c*) | 0.008 | 14.34 | 0.005 | 0.013 | 4.570 | **<.001^***^** |
| **Subtraction** | Indirect | Age → ∆P300 amplitude → RT(path *a × b*) | 0.001 | 0.001 | –0.001 | 0.002 | - | - |
|  | Component | Age → ∆P300 amplitude (path *a*) | 0.134 | 0.302 | –0.191 | 1.006 | 1.352 | .179 |
|  |  | ∆P300 amplitude → RT (path *b*) | 0.103 | 0.001 | –0.001 | 0.002 | 1.139 | .257 |
|  | Direct | Age → RT (path *c’*) | 0.009 | 0.002 | 0.005 | 0.013 | 4.740 | **<.001^***^** |
|  | Total | Age → RT (path *c*) | 0.010 | 0.002 | 0.006 | 0.013 | 4.930 | **<.001^***^** |

*Note.* The values in bold indicate significant effects after applying Bonferroni correction. ∆P300 amplitude was measured in the right paterio-occipital region (P4, P8, O2). All *β* values were standardized. ^***^*p* < .001, ^**^*p* < .01, ^*^*p* < .05.
